# Mutually exclusive spectral biclustering and its applications in cancer subtyping

**DOI:** 10.1101/2022.04.24.489301

**Authors:** Fengrong Liu, Yaning Yang, Xu Steven Xu, Min Yuan

**Affiliations:** Department of Statistics and Finance, University of Science and Technology of China, Hefei, Anhui 230026, China; Genmab US, Inc, Princeton, NJ 08540, USA; Center for Big data in Public Health, Anhui Medical University, Heifei, Anhui 230032

## Abstract

Many soft biclustering algorithms have been developed and applied to various biological and biomedical data analyses. However, until now, few mutually exclusive (hard) biclustering algorithms have been proposed although they can be extremely useful for identify disease or molecular subtypes based on genomic or transcriptomic data. We considered the biclustering problem of expression matrices as a bipartite graph partitioning problem and developed a novel biclustering algorithm, MESBC, based on Dhillon’s spectral method to detect mutually exclusive biclusters. MESBC simultaneously detects relevant features (genes) and corresponding subgroups, and therefore automatically uses the signature features for each subtype to perform the clustering, improving the clustering performance. MESBC could accurately detect the pre-specified biclusters in simulations, and the identified biclusters were highly consistent with the true labels. Particularly, in setting with high noise, MESBC outperformed existing NMF and Dhillon’s method and provided markedly better accuracy. Analysis of two TCGA datasets (LUAD and BRAC cohorts) revealed that MESBC provided similar or more accurate prognostication (i.e., smaller p value) for overall survival in patients with breast and lung cancer, respectively, compared to the existing, gold-standard subtypes for breast (PAM50) and lung cancer (integrative clustering). In the TCGA lung cancer patients, MESBC detected two clinically relevant, rare subtypes that other biclustering or integrative clustering algorithms could not detect. These findings validated our hypothesis that MESBC could improve molecular subtyping in cancer patients and potentially facilitate better individual patient management, risk stratification, patient selection, therapeutic assignments, as well as better understanding gene signatures and molecular pathways for development of novel therapeutic agents.

## Introduction

Biclustering is a powerful machine learning tool that allows for identifying and partitioning related rows and columns of a data matrix, simultaneously. The idea of biclustering was initially proposed by Hartigan [1] and was named as biclustering by Mirkin [2]. Hofmann et al. utilized biclustering for collaborative recommendation systems to identify customers with similar preferences toward certain products [3]. Cheng et al. first applied the biclustering method in gene expression data analysis to identify co-expressed genes [4]. Dhillon proposed a spectral coclustering method in text mining analysis to detect documents with similar words and properties [5]. Since then, biclustering has been applied to a wide range of biological and biomedical data, particularly high-dimensional genomic and transcriptomic data. Comprehensive reviews of different biclustering algorithms can be found in [6–12].

The published biclustering algorithms can be roughly divided into two classes, namely soft biclustering and hard biclustering, according to the overlapping relationship between blocks. Soft biclustering produces overlapping clusters that a data point can belong to multiple clusters, while hard biclustering generates exclusive clusters and each data point belongs to only one cluster. The vast majority of the current biclustering methods fall into the category of soft biclustering, including Cheng and Church [4], CTWC [13], plaid models [14], OPSM[15], xMOTIFs [16], ISA [17], SAMBA [18], QUBIC [19], QUBIC2 [20], and others [21–27]. In biological research, soft biclustering algorithms have been used as tools to perform functional annotation of unclassified genes [28–32], identify different types of modules with interacting molecules [33–42], and gene signatures and pathways detection [43–48].

So far, fewer algorithms have been developed for non-overlapping hard biclustering, particularly for detection of mutually exclusive biclusters on the diagonal of a matrix [5, 49]. Mutually exclusive biclustering can be extremely useful in identifying clinically relevant disease subtypes. The current molecular subtyping is mainly based on traditional one-way clustering techniques such as hierarchical clustering and k-means clustering, which aggregate similar patients based on the expression of all available genes in the gene expression matrix. However, it is known that each disease subtype is usually better characterized by a subgroup of signature genes. Therefore, mutually exclusive biclustering may be able to simultaneously detect relevant genes for each subtype and better stratify the subpopulations for a specific disease. The first biclustering algorithm based on bipartite graph is proposed by Dhillon, who used eigenvectors of the Laplacian matrix of the bipartite graph to identify mutually exclusive biclusters [5]. In addition, non-negative matrix factorization (NMF) is a popular dimensionality reduction and factorization-based biclustering algorithm that can also detect mutually exclusive biclustering patterns [49]. NMF has been proven useful in many cancer subtyping studies due to its easy interpretation and desired performances [50–54].

Spectral clustering method was originally proposed for traditional one-way clustering of samples (rows of data matrix), in which samples are represented as nodes of a weighted graph and similarity measures of sample pairs as adjacency weights. It is widely known that spectrum (eigenvectors) of normalized adjacency matrix or Laplacian matrix exhibits cluster or connectivity property of the graph [55] and can be used to uncover clusters by applying traditional clustering techniques to eigenvectors [56]. Dhillon (2001) proposed the spectral biclustering method by embedding the rows and columns of a data matrix into a bipartite graph [5]. For this derived bipartite graph, application of the aforementioned (one-way) spectral clustering method ends with biclustering of both samples and variables. Kluger et al. (2003) applied similar ideas in biclustering gene expression data [57].

In this paper, we focused on identifying mutually exclusive biclusters using spectral clustering method and proposed to construct a novel biclustering algorithm using principal components (Mutually Exclusive Spectral Biclustering, MESBC) based on spectral graph theory [5, 57]. Simulations showed that MESBC outperformed NMF and Dhillon’s method, providing markedly better accuracy particularly in setting with high noise. That is, MESBC can accurately detect the pre-specified biclusters in simulations, and the biclusters identified by MESBC are highly consistent with the true labels. In real world analysis, MESBC provided more accurate prognostication (i.e., smaller p value) for overall survival in the TCGA LUAD and BRCA cohorts, which potentially can facilitate better individual patient management and risk stratification, as well as better understanding gene signatures and molecular pathways associated with the newly identified subtypes for lung and breast cancer. The rest of paper is organized as follows. In section 2, we formulate the task of biclustering as graph partition. Section 3 gives the simulation results. In section 4, we applied the new proposed biclustering method to two cancer gene expression datasets. Finally, we conclude with a discussion.

## Methods

Suppose we have a nonnegative gene expression matrix *W_m×p_* with rows representing genes and columns representing conditions. The goal of biclustering the gene expression matrix *W* is to partition the genes and conditions into *k* exclusive and exhausted subsets to form *k* biclusters. Denote the row and column labels of *W* be *I* = {*gene*_1_, *gene*_2_,…, *gene_m_*} and *J* = {*cond*_1_, *cond*_2_,…, *cond_p_*} respectively. Suppose *I* and *J* could be divided into *k* disjoint and nonempty subsets {*I*_1_, *I*_2_,…, *I_k_*} with *I*_1_ ∪… ∪ *I_k_* = *I* and {*J*_1_, *J*_2_,…, *J_k_*} with *J*_1_ ∪… ∪ *J_k_* = *J* respectively, then {*I*_1_ × *J*_1_, *I*_2_ × *J*_2_,…, *I_k_ × J_k_*} forms *k* mutually exclusive biclusters of gene expression matrix *W*.

By embeding the gene expression matrix *W* into a weighted bipartite graph, we can formulate the task of biclustering as a graph partition problem. Nodes include all genes and conditions which form the two parts of the bipartite graph *G* with *n* = *m* + *p* nodes. The corresponding *n*×*n* adjacency matrix of bipartite graph *G* is denoted as

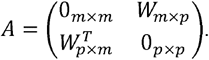

Let 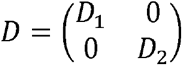 be the degree matrix of *A*, where *D*_1_ and *D*_2_ are diagonal matrices such that 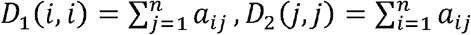. Without of loss of generality, we assume that all the diagonal elements of *D* are positive so that *D* is invertible. Let *B* = *D*^−1^*A* be the random walk normalizd adjacency matrix, then the non-zero eigenvalue of *B* are 1 = *λ*_1_ ≥… ≥ *λ_r_* > 0 > –*λ_r_* ≥… ≥ *λ*_1_ = −1, where *r* – rank(*W*), Suppose 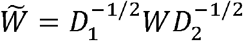 has the singular value decomposition 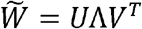, where Λ = *diag*(*λ*_1_,…, *λ_r_*) and *U^T^U* = *V^T^V* = *I_r_*. It can be shown that the eigenvector matrix of *B* corresponding to positive eigenvalues 1 = *λ*_1_, ≥… ≥ *λ_r_* > 0 are 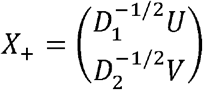 and the principal components are *Y* = *X*_+_Λ, the scaled eigenvectors by eigenvalues.

As the eigenvector for the largest eigenvalue of *B* has the form (*a,a,b,b,b,c,c,…*), Dhillon (2001) argued that biclustering or bipartite partition can be performed based on eigenvectors of the largest eigenvalue [5]. Similar but different from Dhillon’s approach, we propose a mutually exclusive spectral biclustering algorithm (MESBC) for analysis of gene expression data based on the largest *k* principal components of matrix *B* = *D*^−1^*A* instead of eigenvectors of Laplacian matrix *L* = *D* – *A*. As can be observed in simulation studies, the principal components, as scaled eigenvectors, are able to provide better performance in biclustering than that based on the eigenvectors. The proposed biclustering procedure contains the following steps:

**Step 1:** Preprocess the original gene expression matrix *W* to ensure that all elements are non-negative. For example, we can add each element of *W* with the absolute value of the minimum negative number of all elements of *W*, that is, *W* + |*min*(0, *min*(*W*)|. Construct adjacency matrix *A* and degree matrix *D* for the corresponding bipartie graph based on preprocessed non-negative expression matrix *W*. Normalize *W* and perform singular value decomposition, namely, 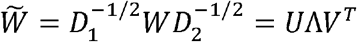.
**Step 2:** Select *k* columns of *U* and *V* corresponding to the top *k* eigenvalues of 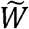 to get matrices *U_k_* = (**u**_1_,…, **u***_k_*) and *V_k_* = (**v**_1_,…, **v***_k_*) respectively. Stack the k*^th^* order principal components 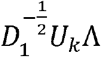 and 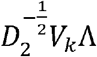 as 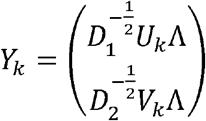.
**Step 3:** Perform k-means algorithm on *Y_k_* to obtain *k* clusters. Identify *k* biclusters according to labels of genes and conditions from k-mean clustering results. The optimal value of *k* is selected that with maximal graph modularity (see the next section for details).

## Simulations

### Parameter settings

We use extensive simulations to compare MESBC with NMF and Dhillon’s method under various scenarios. We assume that there are five biclusters with different sizes, i.e., 20 × 20, 50 × 20, 50 × 20, 30 × 40 and 100 × 50 respectively. Non-bicluster elements (i.e., the background noise) are assumed to be randomly generated from Poisson distribution with mean 1 and elements within the *i*th bicluster are sampled from Poisson distribution with means *λ_i_, i* = 1, 2,…, 5. Parameter *λ_i_* stands for signal-noise ratio and is randomly sampled from the sets [5], {3, 4, 5} and {6, 7, 8, 9} respectively accounting for the low to high signal-to-noise ratios. Each scenario was simulated for 100 replicates. We visualized and compared the biclusters produced by MESBC and the other two clustering approaches with heat-maps. We also evaluated the performance with various internal measures including modularity, silhouette coefficient, ARI, recovery, relevance, the accuracy, precision, recall and F_1_ score.

### Biclusters visualization

Figure 2 showed that when the signal-noise ratio *λ*) and the bicluster size are small, Dhillon’s method and NMF clearly had difficulties to identify the underlying true biclusters, whereas MESBC could still recover the true patterns with high accuracy (Figure 3). The performance of all of the three biclustering algorithms improved as the signal-noise ratio increased. All five true biclusters could be correctly identified with MESBC and NMF when the signal-ratio is greater than 2 (Figure 2b and 2c).

**Figure 1:**
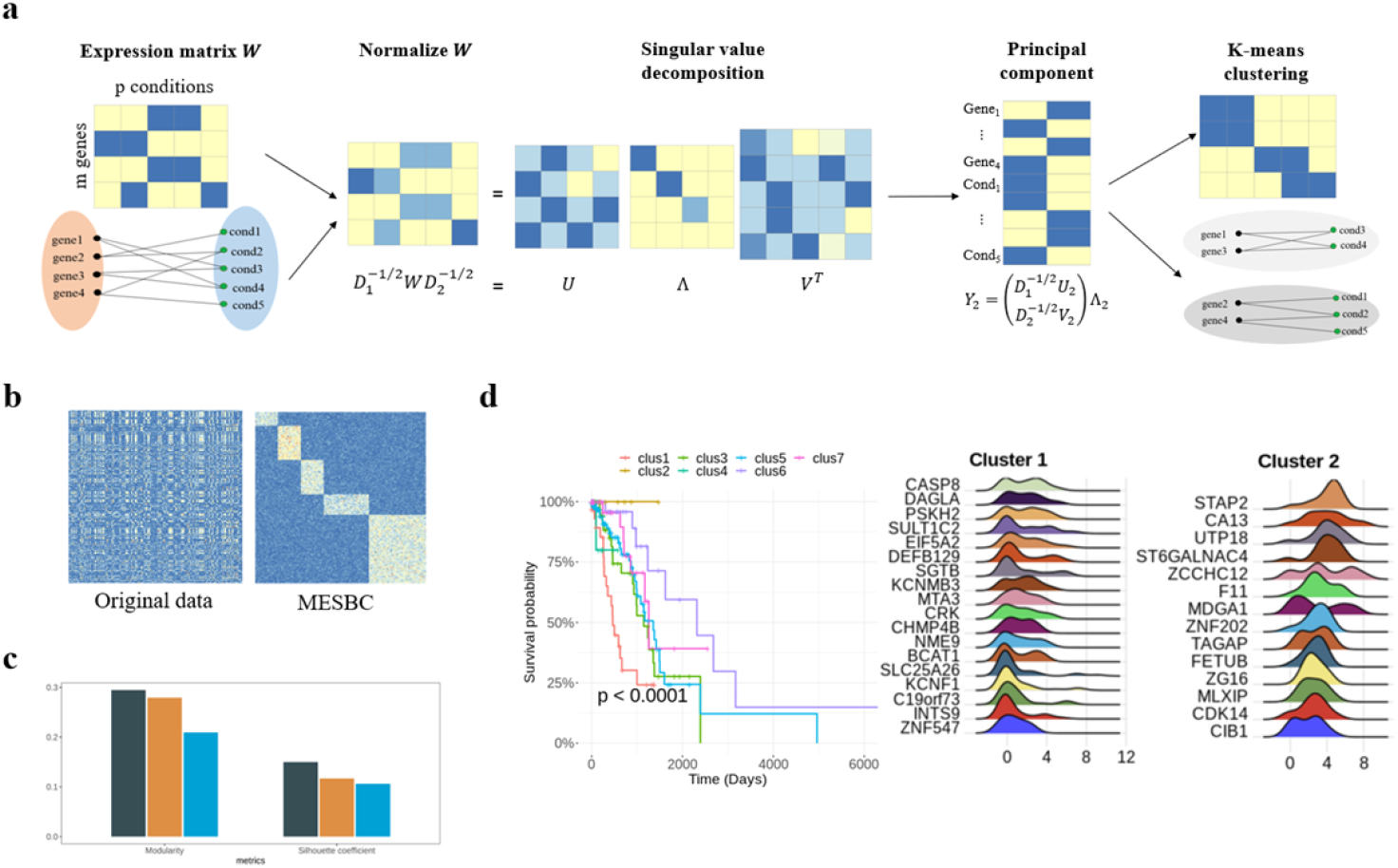
The analysis framework for MESBC of gene expression matrix. (a) Overview of biclustering with MESBC of gene expression matrix with m genes and p conditions (m=4, p=5). (b) Heatmap for original data and adjusted data according to MESBC biclustering results. (c) Quantitative analysis of biclustering results with modularity and silhouette coefficient. For simulation data, there are also internal measures including ARI, recovery, relevance, the accuracy, precision, recall and F1 score. (d) Ridge map and survival analysis of identified subtypes.

**Figure 2:**
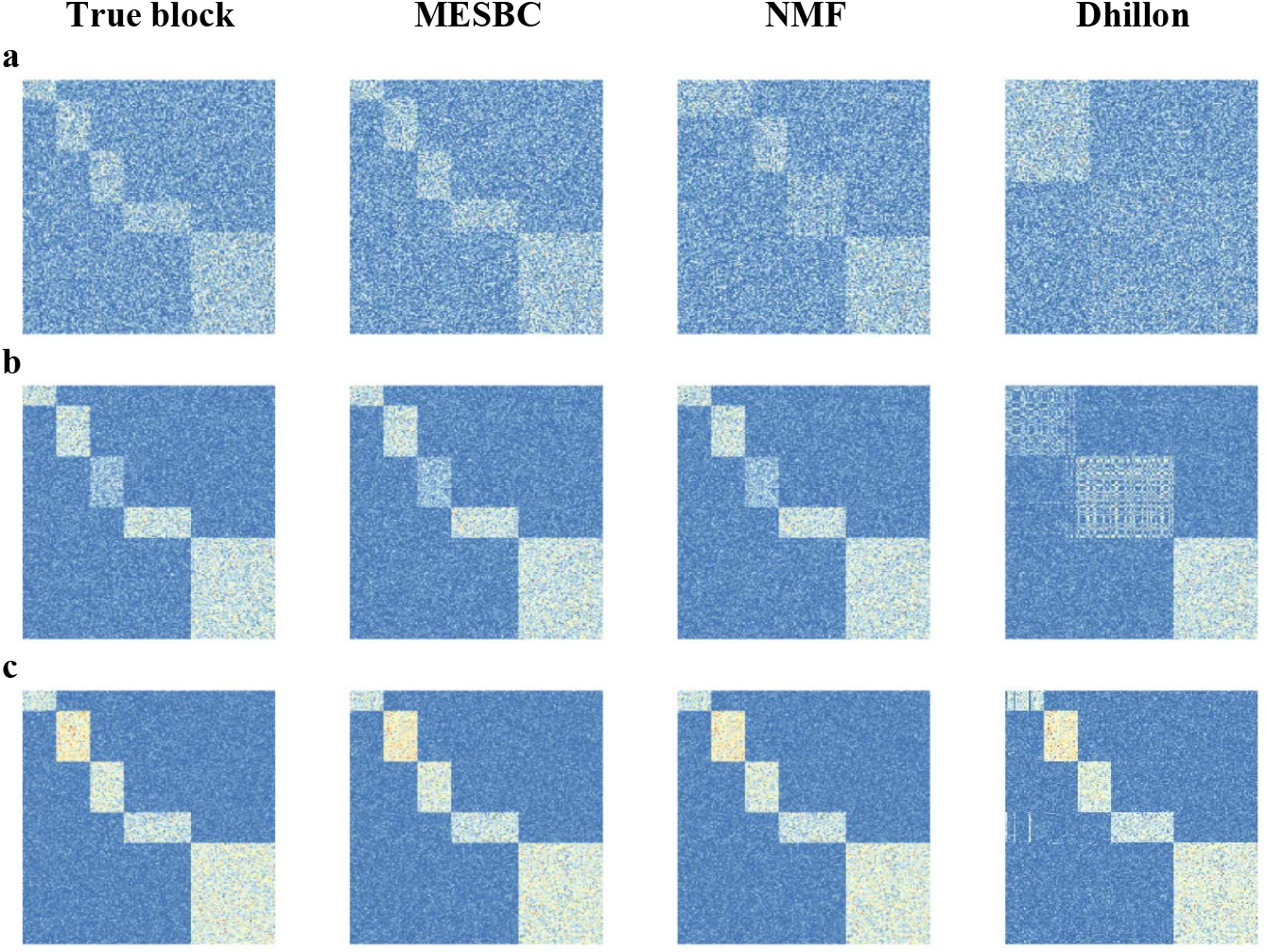
Comparison of biclusters from MESBC, NMF and Dhillon’s method to true blocks using heatmaps at different signal-noise ratios (random sampling from signal-noise ratios = {2} (a), {3, 4, 5} (b) and {6, 7, 8, 9} (c)).

**Figure 3:**
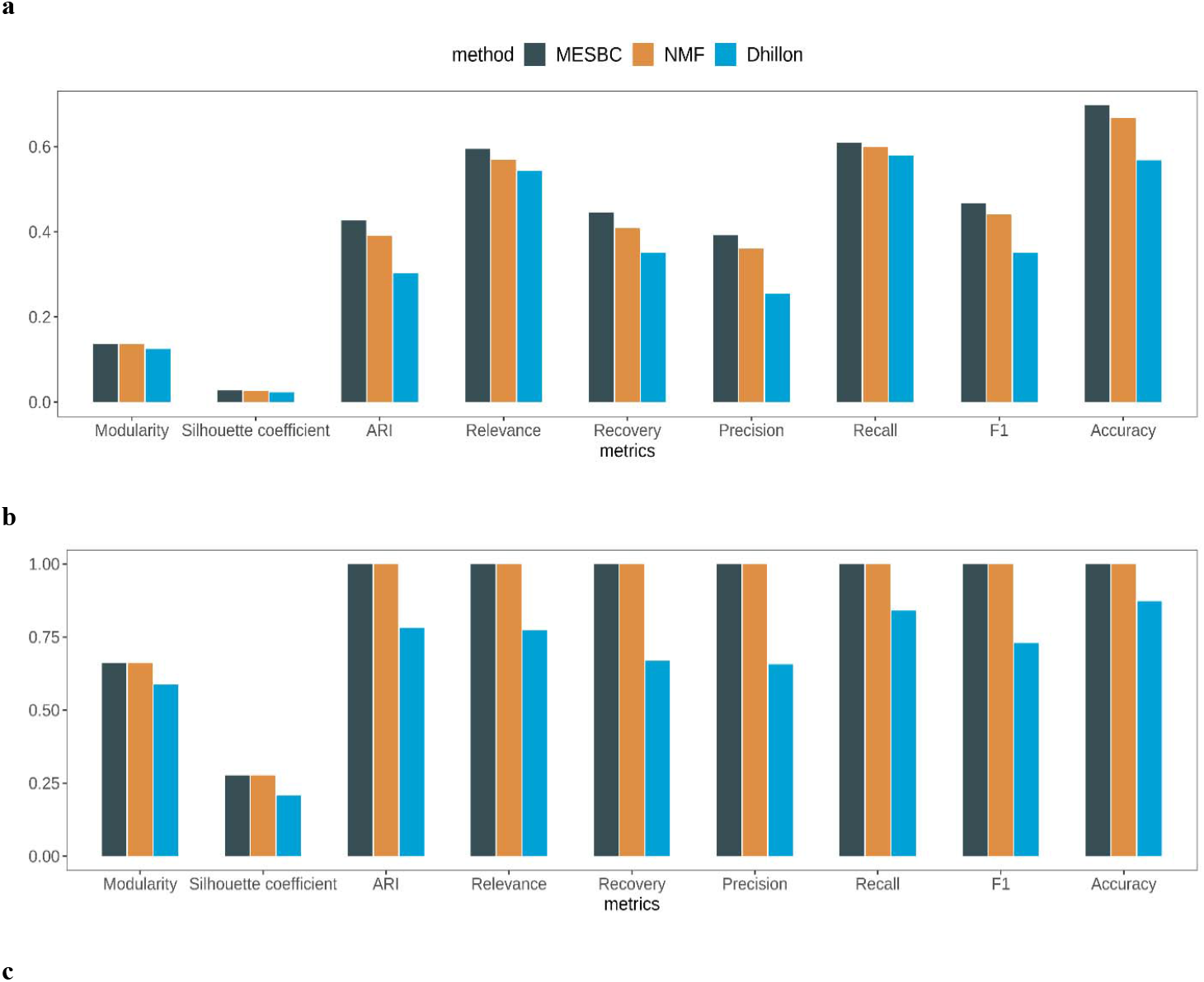
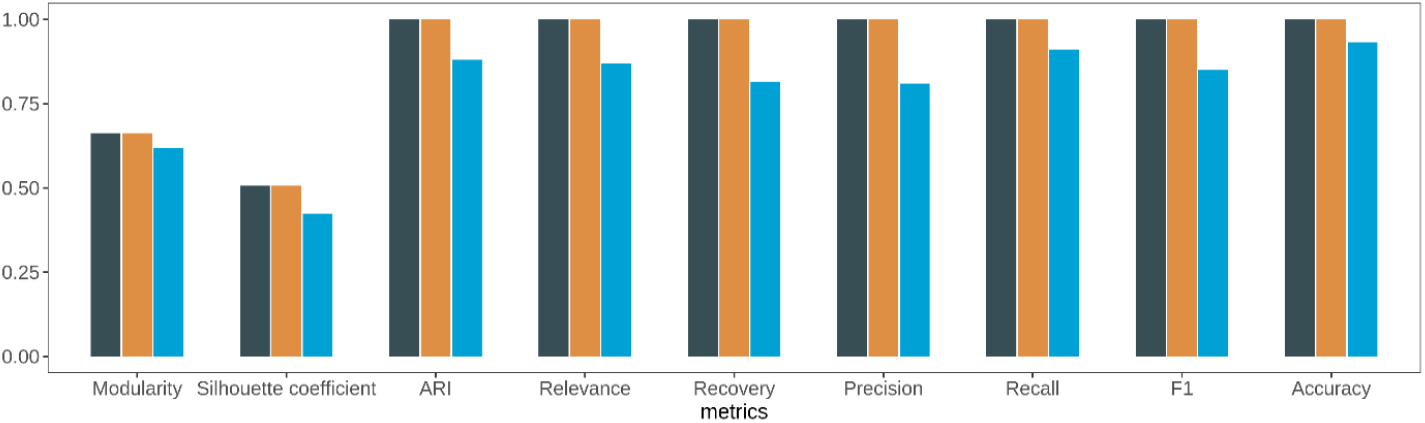
Comparison of the performance among MESBC, NMF and Dhillon’s method at different signal-noise ratios (random sampling from signal-noise ratios = {2} (a), {3, 4, 5} (b) and {6, 7, 8, 9} (c)).

### Clustering metrics

Figure 3 presented the modularity, silhouette coefficient, ARI, relevance, recovery, precision, recall, F1 score and accuracy. Consistent with the heatmaps, the performance of MESBC was clearly the best among the three evaluated algorithms when the signal-noise ratio was small (*λ* = 2, high background noise compared to signal, Figure 3a). At *λ* = 2, although NMF had worse performance than MESBC, it still generally provided better accuracy and separation when compared to Dhillon’s method. When the signal-noise ratios increased to medium (*λ* ranged from 3 to 5) or large (*λ* ranged from 6 to 9), the performance of all the three algorithms improved (Figure 3b and 3c). However, at the medium or high signal-noise ratios, MESBC and NMF appeared to share comparable performance and could correctly identified the true underlying biclusters as ARI approached 1 for both algorithms. Similar to the observations at *λ* = 2, both MESBC and NMF outperformed the Dhillon’s method.

## Real Data Analysis

### Dataset

The Cancer Genome Atlas (TCGA) lung adenocarcinoma (LUAD) and breast cancer (BRCA) count data was downloaded from legacy database using GDC Application Programming Interface (API) by ‘TCGAbiolinks’ package [58]. Feature counts for each cell were divided by the total counts of that cell and multiplied by 10,000, and we applied a natural log transformation using log(1 + *x*). In the bulk gene expression datasets, most genes are lowly expressed and do not vary to a larger extent. Therefore, we filtered out the top 100 genes for this analysis. Cancer subtype labels were obtained from PanCancerAtlas_subtypes in ‘CancerSubtypes’ package [59]. Survival time and status for patients were downloaded from the National Cancer Institute (NCI) Genomic Data Commons (GDC) (https://portal.gdc.cancer.gov/). After excluding samples without label or survival information, we used gene expression data with 268 (LUAD) and 1204 (BRCA) patients for biclustering analysis, and compared MESBC with Dhillon’s method and NMF algorithm which is widely applied to subtype identification [49].

### Optimal number of biclusters

The optimal number of biclusters was determined by modularity, which measures the degree of connectivity within clusters relative to between clusters. [60–62]. In this study, we first constructed a shared nearest-neighbor (SNN) [63] graph considering ten shared nearest neighbors using the first fifty principal components of *Y* = *X*_+_Λ by buildSNNGraph in R package ‘scran’. The modularity was then computed based on the constructed network. After running the algorithm with the number of biclusters *k* from 2 to 10, the optimal value was defined as the *k* value corresponding to maximum modularity.

Figure 4 shows that, for both LUAD and BRCA datasets, MESBC clearly provided higher modularity across the entire range of *k* (from 2 to 10) compared to Dhillon and NMF, indicating that the biclusters from MESBC had the strongest network community structure among the three algorithms. For LUAD, the modularity of MESBC increased from *k* = 2 to 4, leveled off from *k* = 4 to 7, and then slightly decreased afterwards (*k* > 7). A k value of 7 provided the highest modularity. The modularity of NMF peaked at *k* = 4, while Dhillon’s method showed high fluctuations at low levels of modularity. A *k* value of 9 appears to produce the highest modularity for Dhillon’s method. For BRCA, the modularity of MESBC peaked at *k* = 6, whereas NMF and Dhillon’s method reached the highest modularity at *k* = 4 and *k* = 5, respectively.

**Figure 4.**
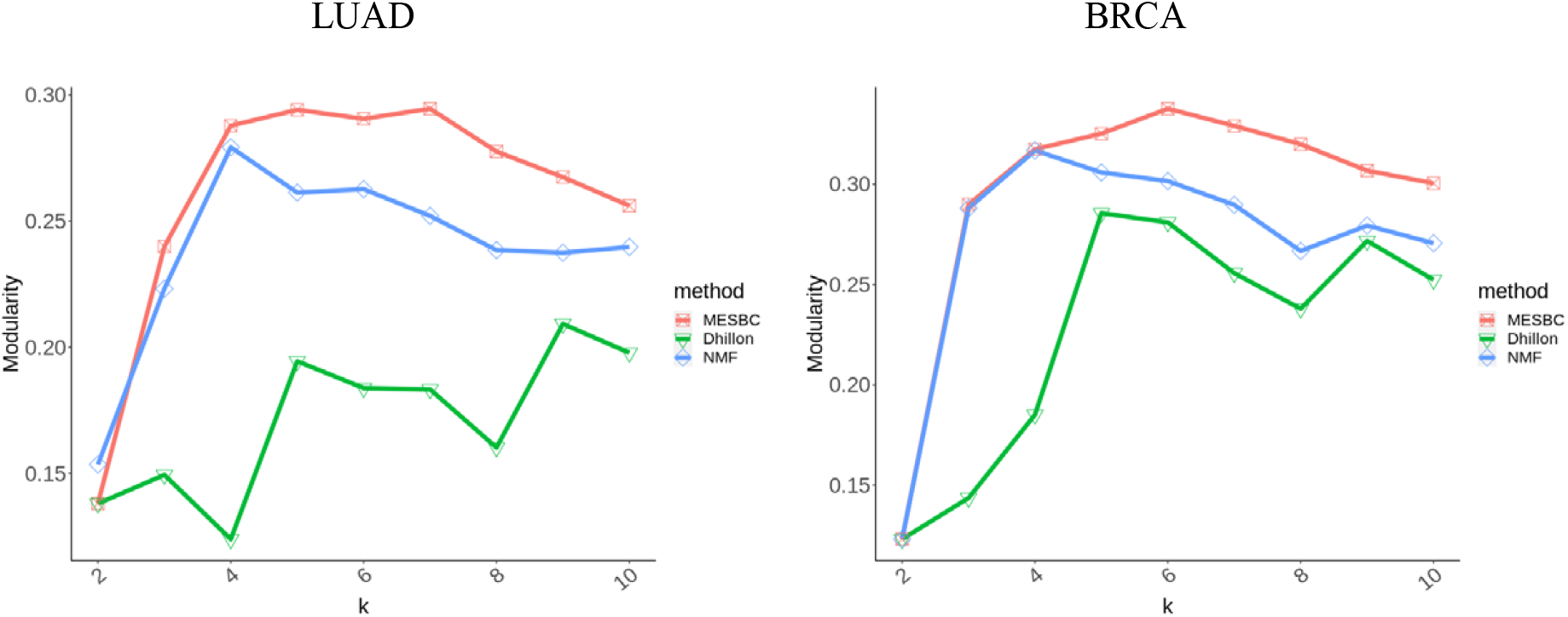
Modularity with the number of biclusters *k* from 2 to 10 for MESBC, NMF and Dhillon’s method in LUAD and BRCA data.

### Biclustering pattern of gene expression

Compared to conventional clustering, mutually exclusive biclustering (MESBC, Dhillon, and NMF) can directly identify the corresponding gene signatures for each cluster (subtype). Figure 5 illustrates biclusters using heatmaps, where the rows (patients) and columns (genes) were arranged according to the number of the cluster. At the optimal number of biclusters, all the three biclustering methods can dissect the unstructured data and discovered apparent modules and patterns of both the LUAD (Figure 5a) and BRCA (Figure 5b) expression data.

**Figure 5.**
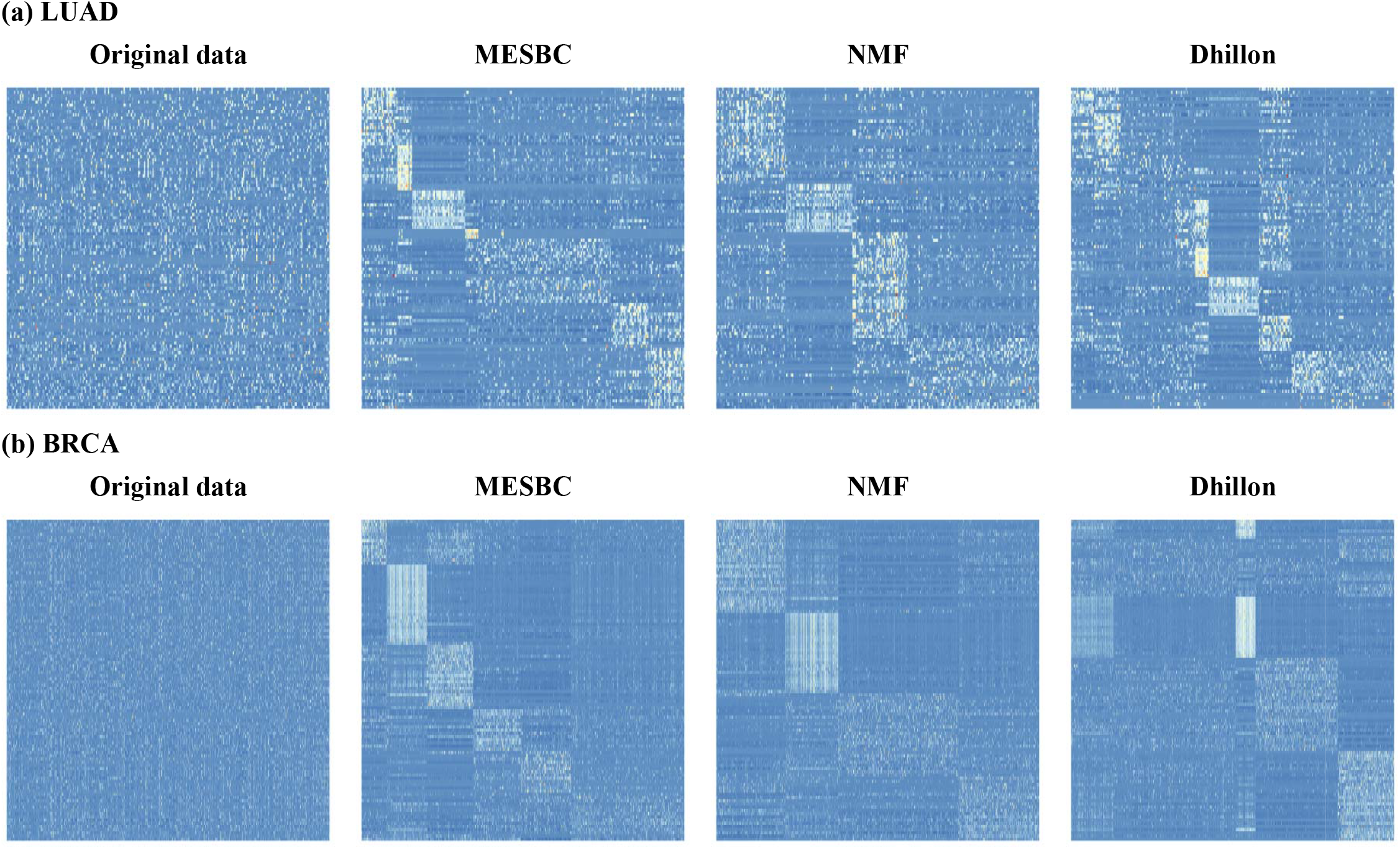
Heatmap for MESBC, NMF and Dhillon’s method. The number of biclusters k is selected according to maximum modularity.

With regard to LUAD dataset (Figure 5a), in contrast to Dhillon generating excessive clusters (*k*=9) with some unobvious modules and NMF identifying fewer clusters (*k*=4) with weak internal connections, MESBC detected 7 biclusters with more evident internal consistency and external heterogeneity. For the BRCA dataset, the pattern of the biclusters detected by MESBC was clearly more obvious than that detected by NMF and Dhillon’s method (Figure 5b). Particularly for Dhillon’s method, there appears no clearly pattern of the biclusters identified along the diagonal of the gene expression matrix.

### Quantitative analysis

Modularity and silhouette coefficient [64] were used to quantify the performance of the three different biclustering algorithms (Figure 6). For LUAD, MESBC clearly provided the highest modularity and silhouette coefficient among the three algorithms, while Dhillon yielded the lowest modularity and silhouette coefficient. Similarly, for BRCA dataset, MESBC consistently produced the highest modularity although NMF appears to have slightly higher silhouette coefficient than MESBC. Consistent with findings from LUAD, Dhillon produced the worst performance based on BRAC. In general, MESBC outperformed NMF and Dhillon’s method in terms of tightness and separation in both LUAD and BRAC.

**Figure 6.**
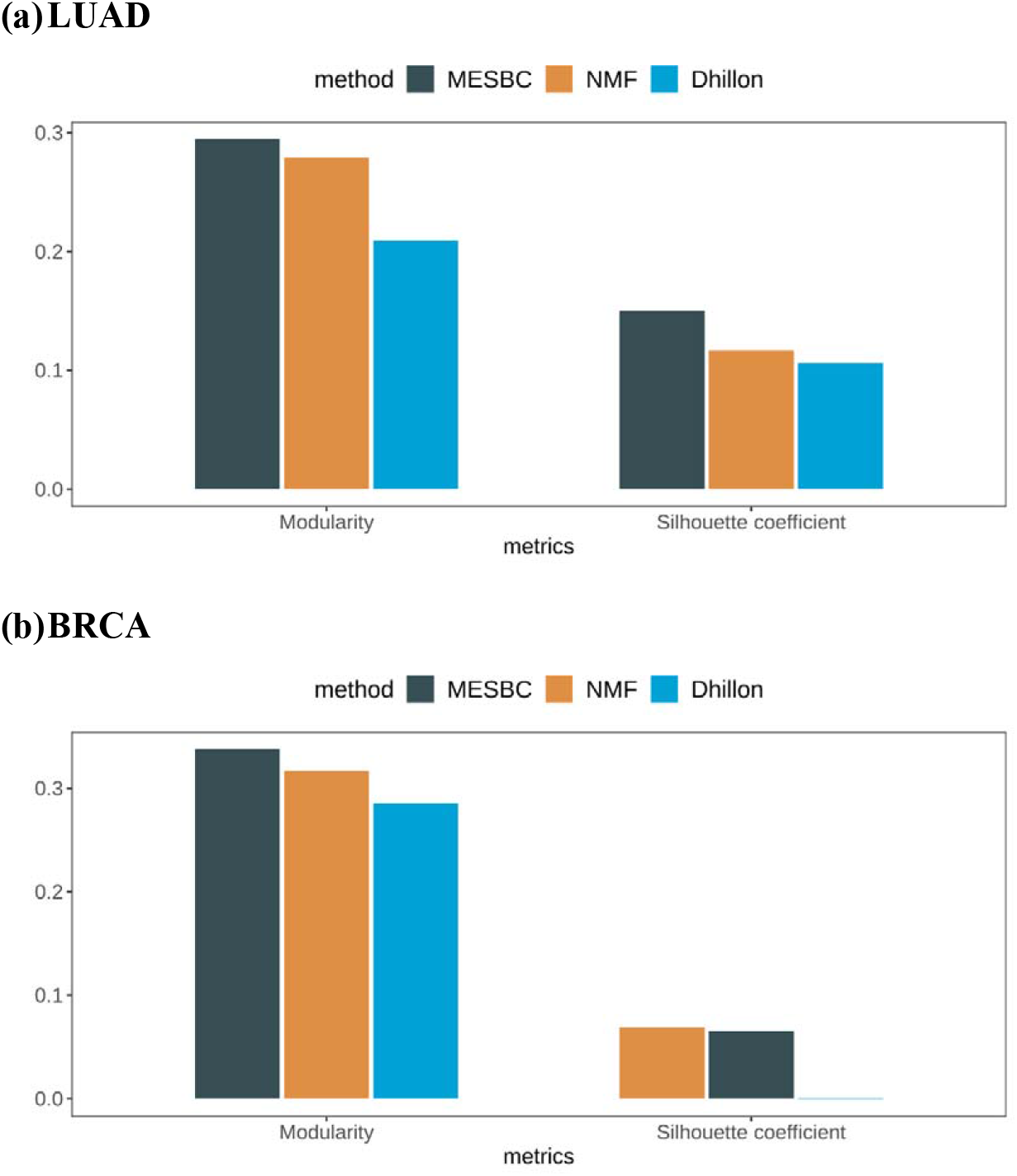
Bar Plot of modularity and silhouette coefficient and ARI for MESBC, NMF and Dhillon’s method.

### Risk stratification

We conducted survival analysis using Kaplan-Meier method according to the subtypes identified by the three biclustering algorithms and compared the results to published TCGA subtypes [65]. Survival curves were derived from Kaplan-Meier estimates and the log-rank test was used to evaluate the statistical significance of biclusters (Figure 7). For LUAD, the subtypes (biclusters) identified by MESBC demonstrates very distinct survival patterns with highly significant a p value (< 0.0001). In contrast, NMF and Dhillon were only marginally significant (p ~ 0.02). Similarly, the published TCGA subtypes had a p value around 0.02, much less significant compared to that based on MESBC subtypes. Therefore, molecular subtyping based on MESBC could provide more accurate prognostication and better optimized risk stratification for LUAD patients. The more significant result produced by MESBC can be probably attribute to Clusters 1 and 2. The patients in Cluster 1 had very aggressive disease progression leading to large number of deaths within short time of period whereas the patients in Cluster 2 appears to have extremely slow growing disease and no patients died in this group within the follow-up period. More importantly, these two groups of patients were not detected either by the Integrative Clustering (iClust from TCGA using DNA copy number, DNA methylation, and mRNA expression data) [66], NMF, or Dhillon methods. The Sankey diagram in Figure 8 demonstrates the relationship between the published TCGA subtypes and the MESBC subtypes. The subtypes of MESBC have no approximate correspondence with TCGA subgroups, indicating that the clustering mechanism of MESBC is not a simple repetition of existing methods. More specifically, about 60% MESBC-based Cluster 1 patients were consistent with iClust6, which was the subtype with most aggressive disease identified by integrative clustering [67]. However, Cluster 1 of MESBC also consisted of patients from iClust1-4. Further examination of the MESBC-detected signature genes for each subtype, the top three over-expressed gene in Cluster 1 were CASP8, DAGLA, and PSKH2, while the top three genes in Cluster 2 were STAP2, UTP18, and CA13 (Figure 9).

**Figure 7.**
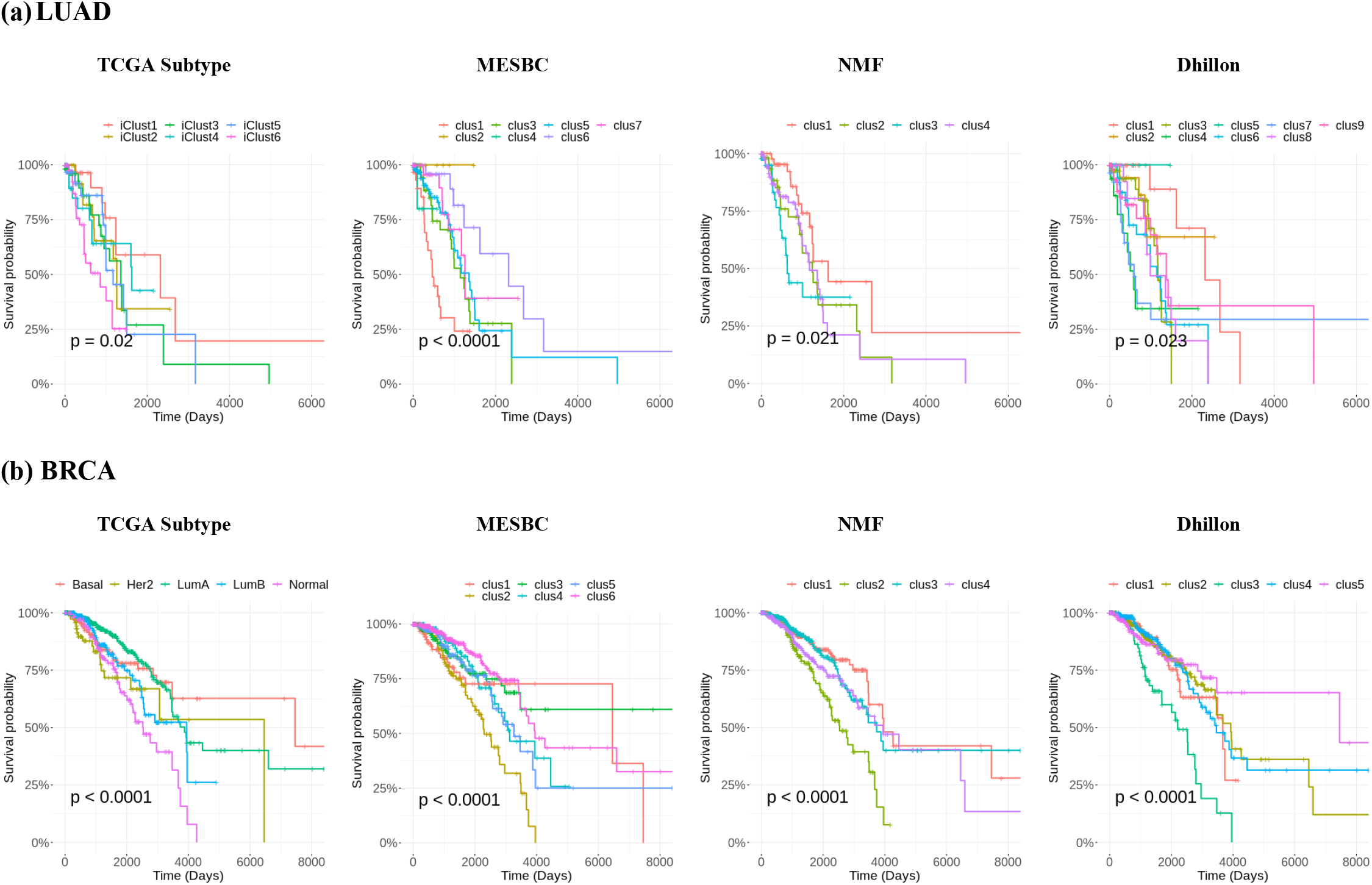
Kaplan Meier survivals plot for MESBC, NMF and Dhillon’s method.

**Figure 8.**
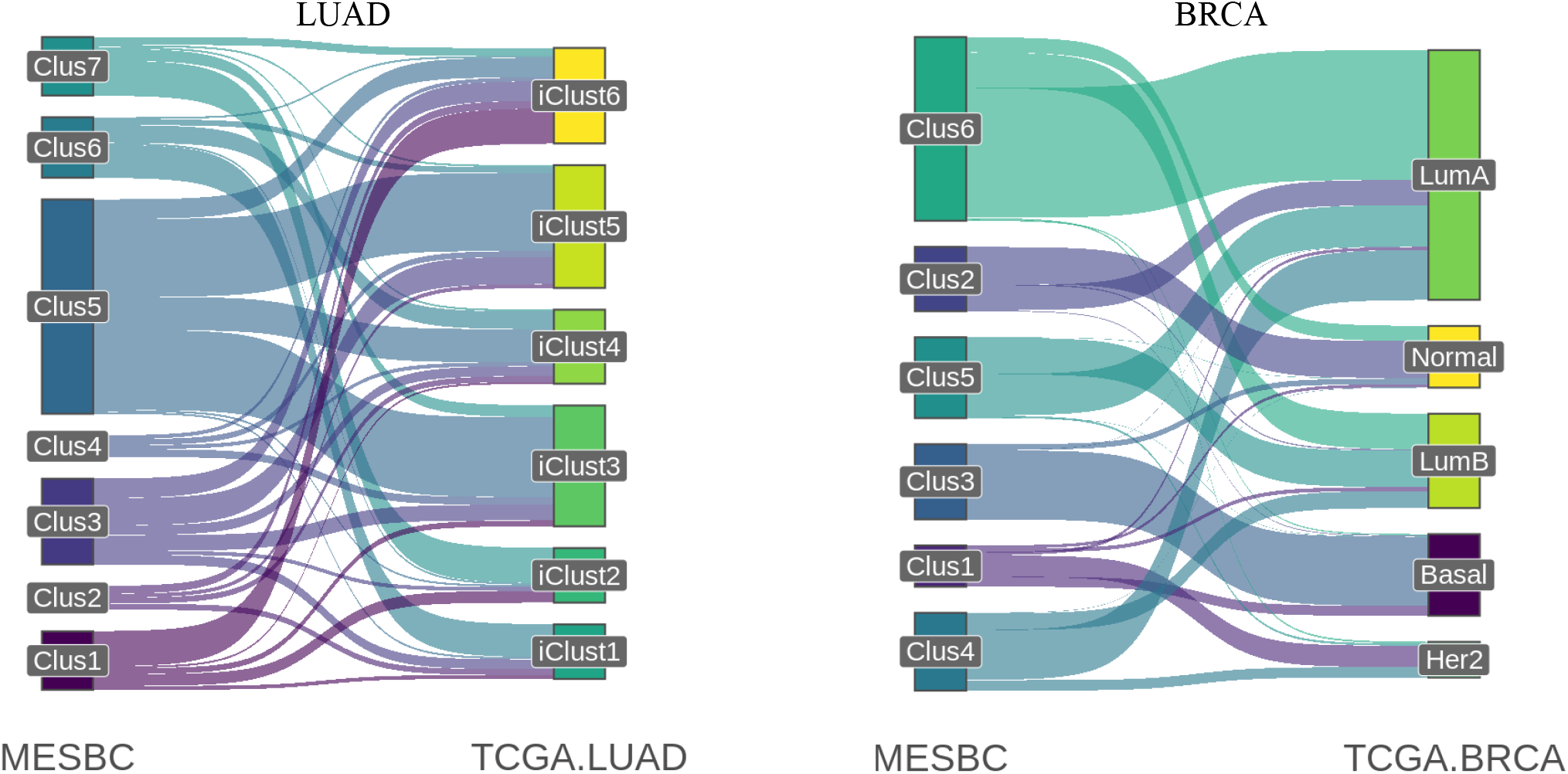
Sankey plots comparing clusters from MESBC with published TCGA subtypes.

**Figure 9.**
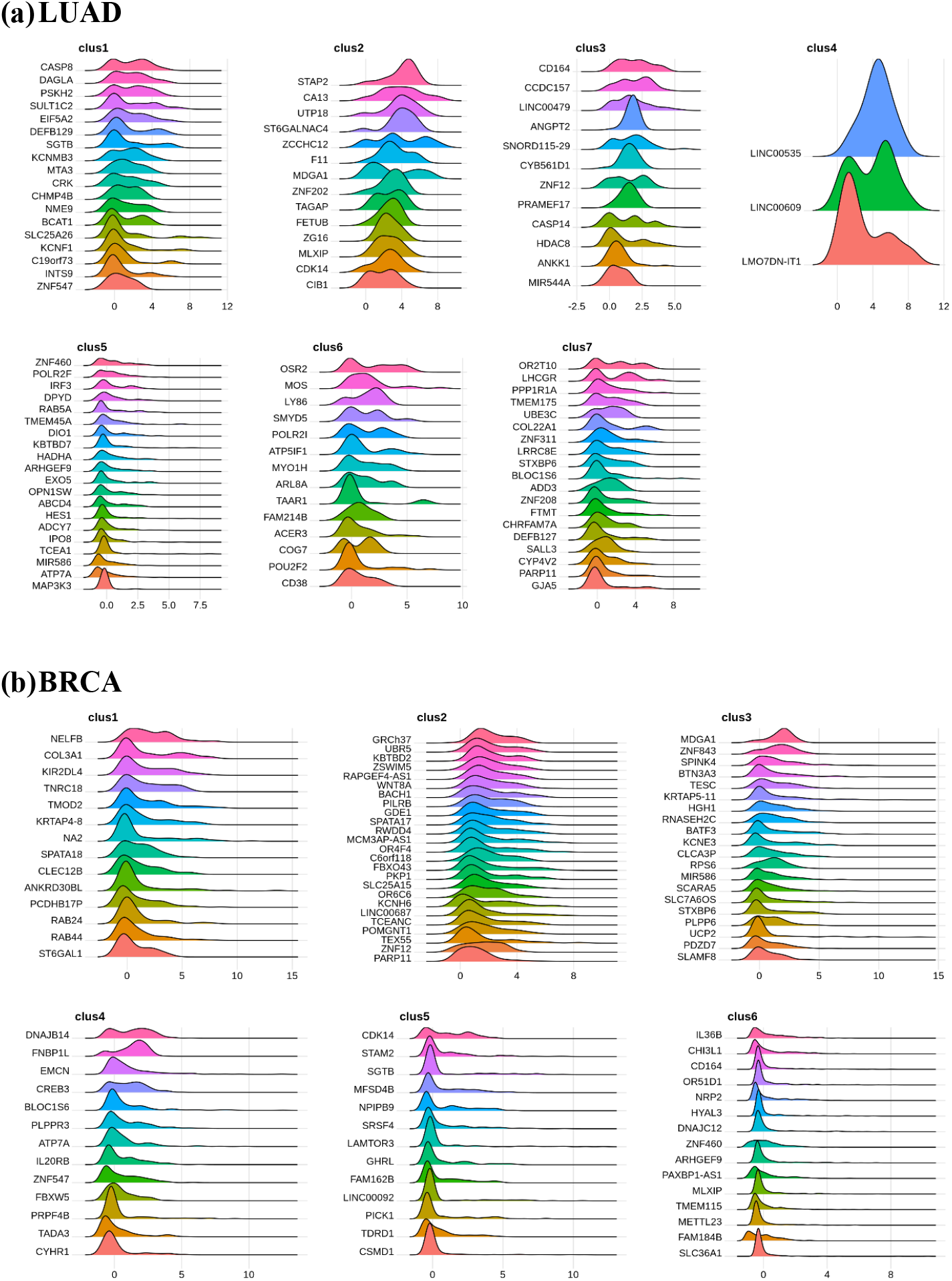
Distribution of expression of each gene in patients from same bicluster (a) LUAD; (b) BRCA.

For BRCA, MESBC provided similar risk classification compared to NMF and Dhillon, as significant association between survival and the subtypes identified by all the three algorithms was observed (Figure 7). In addition, compared to LUAD, MESBC clustering showed greater similarity with PAM50 (Figure 8) for BRCA. The Cluster 3 of MESBC was similar to the Basal subtype based on PAM50. MESBC-based Cluster 2 mainly consisted of Luminal A- and Normal-like patients, while Cluster 5 were mainly Luminal A and B patients. This result is consistent with the fact that both MESBC and PAM50 shared similar prediction performance for overall survival of BRCA patients. The ability to consistently identify clinically relevant subtypes in both LUAD and BRCA cancer patients demonstrated the improved robustness of MESBC compared to other existing subtyping algorithms.

## Discussion

A large number of soft biclustering algorithms have been developed and applied to various biological and biomedical data, including functional annotation of unclassified genes [28–32], detection of different types of modules with interacting molecules [33–42], and identification of gene signatures and pathways [43–48]. However, until now, very few biclustering algorithms have been proposed to identify mutually exclusive biclustering patterns, which can be utilized for molecular subtyping based on genomic or transcriptomic data. In this paper, we developed MESBC, a novel biclustering algorithm based on spectral graph theory to detect mutually exclusive biclusters. Our simulations revealed that MESBC provided superior accuracy to detect pre-specified biclusters compared to NMF and Dhillon’s method, particularly in presence of large degree of noise in the data. In addition, real world analysis of TCGA datasets (LUAD and BRAC) suggest that MESBC could provide similar or more accurate prognostication (i.e., smaller p value) for overall survival in patients with breast and lung cancer, respectively, compared to the existing, gold-standard subtypes for breast (PAM50) and lung cancer (Integrative clustering [66]).

MESBC simultaneously detects relevant genes and corresponding subgroups, and therefore automatically uses the signature genes for each subtype to perform the clustering, improving the clustering performance (i.e., the tightness within each cluster and separation between clusters) and potentially accuracy for risk stratification. On the other hand, classical one-way clustering techniques (e.g., hierarchical or k-means clustering, etc) use global features i.e., all the available genes in the dataset to group patients without taking into account the real, relevant genes for individual clusters, and therefore potentially introduce more noise while performing the clustering. Usually, for classical clustering techniques, one additional step such as differential gene expression, is required to identify signature genes for each cluster, whereas MESBC identifies the subtypes and signature genes at the same time.

It appears that MESBC may be able to detect clinically relevant, rare subtypes that other biclustering (NMF and Dhillon) or integrative clustering (using different modality of data; such as DNA copy number, DNA methylation, and mRNA expression data for LUAD [66]) could not detect. In LUAD, MESBC identify a group of patients (Cluster 2) that appear to have extremely slow disease progression, whereas Cluster 1 identified by MESBC seems to be hyper progressors that have much more aggressive disease than the subtypes identified by the integrative LUAD subtypes (iCluster [67]). Particularly, with the signature genes detected by MESBC (e.g., CASP8, DAGLA, and PSKH2 for Cluster 1 and STAP2, UTP18, and CA13 for Cluster 2), MESBC can potentially provide better optimized individual patient management and/or risk stratification via more accurate prognostication, improved therapeutic assignments, better patient selection and enrollment in clinical trials, and more importantly, development of novel therapeutic agents, etc.

## Acknowledgement

This work was partially supported by the Natural Science Foundation of Anhui Province (No. 2008085MA09) and the National Science Foundation of China (NSFC), Grant No. 11671375.

